# Imaging the impact of rotifer consumption on bacterial behaviors in the zebrafish gut

**DOI:** 10.64898/2026.01.26.701418

**Authors:** Susana Márquez Rosales, Piyush Amitabh, Emily M. Olmstead, Emily P. R. Avey, Elena S. Wall, Lizett Ortiz de Ora, Travis J. Wiles, Raghuveer Parthasarathy

## Abstract

The gut microbiota influence many aspects of their host’s health and physiology including the digestion of food, and food intake in turn influences the composition of the gut microbiome. However, the ways in which food can alter the behavior of intestinal bacteria remain largely unknown, due in large part to the difficulty of assessing behavior *in situ*. Larval zebrafish provide a model for addressing this gap because of their optical transparency and their ability to be prepared germ-free and then associated with specific microbial species. Using light sheet fluorescence microscopy to visualize bacteria inside the intestines of live zebrafish larvae, we examine the properties of two commensal strains with markedly different physical characteristics. One is a zebrafish-commensal *Enterobacter* species that forms large aggregates in unfed larvae, and the other is a pathobiont *Vibrio* species that is motile and planktonic. Following host consumption of rotifers, a common food, *Enterobacter* clusters disintegrate into motile individuals. *Vibrio* remains planktonic in fed larvae but decreases the activity of its Type VI Secretion System, leading to a strong decrease in damage to host tissue. Our results reveal that feeding can have major impacts on bacterial behavior that should be considered in models of normal gut microbiome dynamics as well as pathogenesis.

## Introduction

Diet significantly shapes the composition of the gut microbiota. Different nutrients influence the microbiota directly by altering substrate availability and indirectly via the host, for example increasing inflammation or changing intestinal permeability [1, 2]. Many studies have quantified how diet alters bacterial composition and diversity [3]. For example, fiber-rich diets increase the availability of microbiota-accessible carbohydrates, which promotes greater microbial diversity and supports the growth of beneficial species such as *Bifidobacterium* [4]. In contrast, high-fat and high-sugar diets shift the microbiota toward dysbiotic, inflammatory states [5, 6]. Diet can also influence microbiota function by altering bacterial gene expression. For example, consumption of meat rapidly leads to increased expression of bacterial genes for amino acid catabolism and other protein digestion tasks, while plant-based diets induce increased carbohydrate-digestion activity [7]. Diet-induced shifts can directly impact the host; in mice, a shortage of dietary fiber induces gut microbes to degrade colonic mucus, facilitating epithelial access for pathogens [8]. In general, however, diet-induced changes in gut bacterial behaviors, especially behaviors related to physical characteristics such as motility and aggregation, remain challenging to study in animals due to their lack of optical accessibility. The variety of ways that behaviors might be altered, therefore, is likely underappreciated, limiting our ability to predict the impact of various diets on the intestinal microbiome.

Zebrafish are a widely used vertebrate animal model for many developmental and physiological processes in large part because of their transparency at young ages. In addition, zebrafish can be prepared sterile (germ-free), after which they can be introduced to specific bacterial strains. As a consequence of these attributes, zebrafish are increasingly used to investigate host-microbe interactions [9]. A collection of zebrafish-isolate strains native to the gut microbiota, engineered to express fluorescent proteins [10], together with light sheet fluorescent microscopy (LSFM), an imaging method that provides rapid, three-dimensional, cell-resolved imaging capable of spanning the entire gut, enables detailed observation of bacterial behavior and spatial organization within live animals [11–13]. In this collection of zebrafish symbiotic bacteria, two strains stand out for their contrasting behaviors in the larval gut: *Enterobacter* ZOR0014 and *Vibrio* ZWU0020. When the sole species present, the *Enterobacter* strain forms dense aggregates in the midgut [13]. In contrast, *Vibrio* ZWU0020 is mainly planktonic, highly motile, and is dispersed throughout the gut. This strain carries a Type VI Secretion System (T6SS) that damages host cells at the posterior opening of the gut (vent) and drives strong gut contractions as a result of immune cell activity [14]. *Vibrio* ZWU0020 is genetically very similar to the human pathogen *Vibrio cholerae*, which also wields a T6SS capable of driving strong intestinal contractions [15]. We have previously demonstrated that each of these strains, *Enterobacter* ZOR0014 and *Vibrio* ZWU0020, can be induced to change its behavior. *Enterobacter* dis-aggregates if the gut is later colonized by a mutant of a zebrafish-isolate *Aeromonas* strain; the dispersed *Enterobacter* individuals are non-motile and fail to persist in the gut [16]. Low doses of the antibiotic ciprofloxacin induce *Vibrio* filamentation, shifting it from a planktonic to an aggregated state that makes it more vulnerable to peristalsis and expulsion from the intestine [17].

So far, nearly all studies have used unfed larvae because feeding while maintaining gnotobiotic conditions is challenging, as larvae grow well when fed live prey, which bring their own microbes. As a result, the impact of food on native bacteria in the zebrafish gut remains largely unknown. In previous work, we developed a protocol to reduce the bacterial load in rotifers, a commonly used live food, by irradiating them with ultraviolet light. This procedure enables us to maintain healthy larvae under near-gnotobiotic conditions [18].

In this study, we use UV-treated rotifers to investigate how food ingestion can influence specific bacterial strains within the zebrafish gut, focusing on the above-mentioned *Enterobacter* ZOR0014 and *Vibrio* ZWU0020, hereafter referred to as *Enterobacter* and *Vibrio* for brevity. We show that rotifer ingestion modifies bacterial behavior in distinct and unexpected ways: *Enterobacter* shifts its spatial organization from non-motile aggregates to planktonic motile cells, and *Vibrio* reduces its T6SS activity, leading to less host tissue damage.

## Results

As noted above, we used recently developed techniques to obtain nearly sterile rotifers using ultraviolet irradiation (Materials and Methods). Because larval zebrafish hunt live rotifers, the time and amount of their consumption is variable. We first determined whether we could detect through imaging the presence of rotifers in the intestines of individual larvae, to establish that feeding had occurred. Zebrafish were derived germ-free using established methods [19] and then at day 5 post-fertilization (dpf) were allowed to hunt UV-irradiated rotifers for around 20 hours. Larvae were imaged the next day (6 dpf) using brightfield and light sheet fluorescence microscopy (LSFM) (Materials and Methods). Rotifers are roughly 100 to 300 *μ*m in extent, the same order of magnitude as the larval gut diameter, so we anticipated they would be visible. Bright-field images confirmed our expectation: fed larvae showed a noticeably larger and darker gut compared to unfed larvae (Fig. 1B–C).

**Figure 1.**
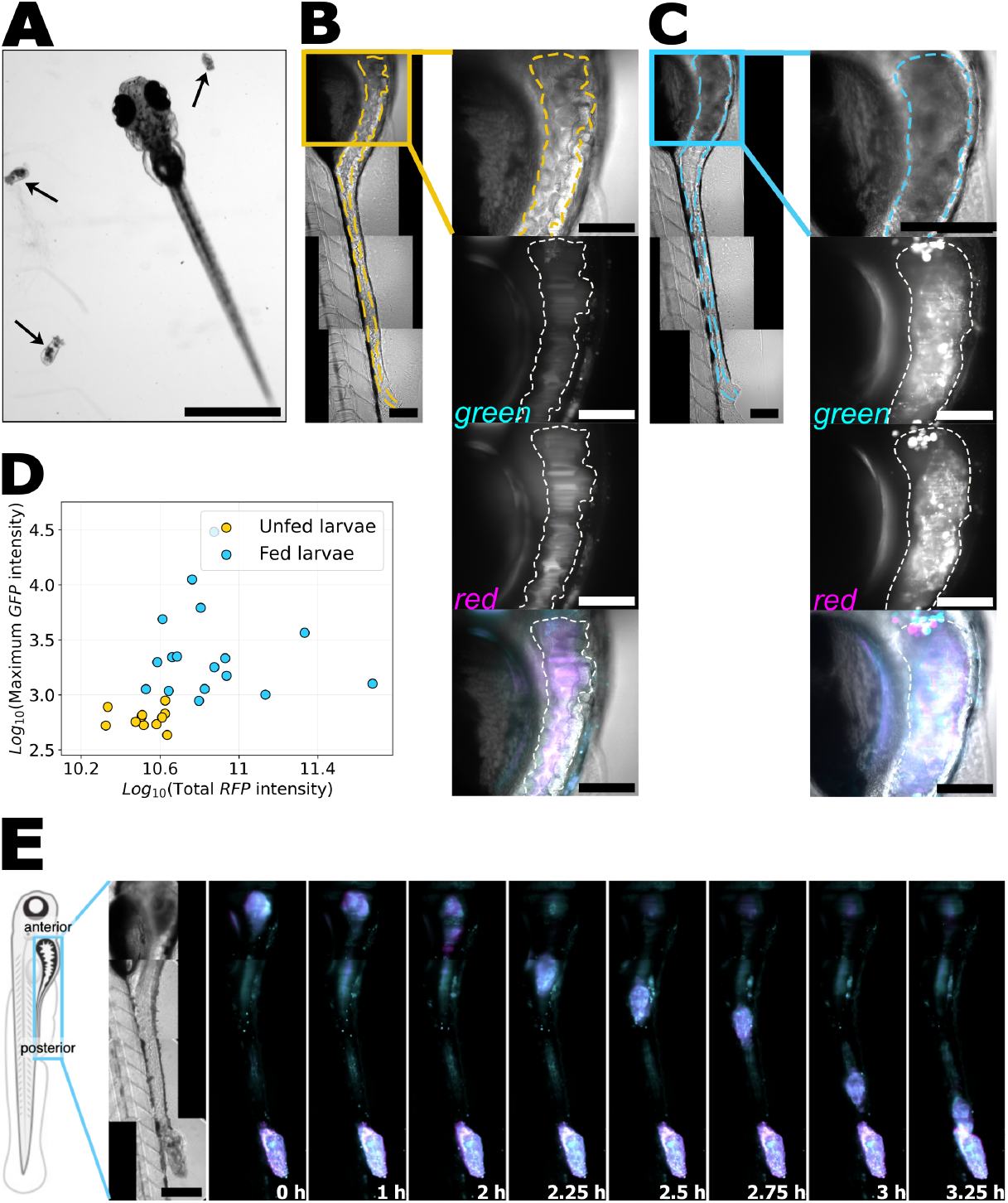
**A** A Zebrafish larva at 5 days post-fertilization (dpf) alongside rotifers (arrows). Scale bar: 1 mm. **B** The gut and surrounding regions of a germ-free, unfed larva at 6 dpf. The dashed line indicates the approximate intestinal boundary. The yellow box includes the anterior gut region, shown in the zoomed-in panels. From top to bottom: bright field, autofluorescence in single optical planes using blue/green excitation/emission wavelengths (excitation 488 nm, emission 510-550 nm, denoted ‘green’), autofluorescence using green/red excitation/emission wavelengths (excitation 561 nm, emission 590-650 nm, denoted ‘red’), and a merged composite image (gray: bright field, cyan: green autofluorescence, magenta: red autofluorescence). **C** Gut of a germ-free 6 dpf larva fed with rotifers, with a rotifer in the anterior gut. Zoomed-in panels include bright field, green and red autofluorescence, and a composite image as in (B). Scale bar: 100 *μ*m. **D** Comparison of gut autofluorescence intensity in unfed (yellow circles) and rotifer-fed (blue circles) larvae. Each dot represents one larval gut. **E** From left to right: schematic of a larval zebrafish with the same orientation as in the following panels; bright field image of a fed larva at the initial imaging time; composite images (cyan : green autofluorescence, magenta: red autofluorescence) from a series of time points showing rotifer transit. Time intervals are not uniform. Scale bar: 200 *μ*m.

We used LSFM to record fluorescence in two channels: one with excitation at 488 nm and emission at 510-550 nm (green), and another with excitation at 561 nm and emission at 590-650 nm (red). Because neither rotifers nor larvae expressed fluorescent proteins, signals in green and red channels resulted solely from autofluorescence. We found that the autofluorescence intensity in both channels was higher in fed than in unfed larvae and showed a crumpled morphology in fed larvae (Fig. 1C, Supplemental figure and video). Quantifying either the maximum or the total autofluorescence intensity provides an objective indication that the feeding state of the fish can be assessed by imaging. Plotting the maximum autofluorescence intensity in the green channel and the total autofluorescence intensity in the red channel shows clear discrimination between fed and unfed animals (Fig. 1D, Materials and Methods), indicating that the feeding state of larval zebrafish is resolvable through imaging (Fig. 1D).

To assess the timescale of rotifer transit from the anterior to the posterior gut, we imaged larvae every 15 minutes for 8 hours (Materials and Methods). We observed the rotifers were retained in the larger-diameter anterior “bulb” region for a few hours. Then, rotifers rapidly transitted to the posterior, typically reaching the the vent in less than 1 hour. In the example shown in Fig. 1E, a rotifer stayed in the anterior for 2 hours. In the example shown in Supplemental Video 3, a rotifer remained in the anterior for more than 5 hours before descending. Overall, the timescale for the presence of rotifers in the gut is on the order of hours.

We next examined how rotifer ingestion influences gut bacteria, focusing on two model species, each labeled with constitutively expressed green fluorescence protein (GFP). We first consider the *Enterobacter* strain that, as noted above, forms large aggregates in unfed fish (Fig. 2A) [13]. We inoculated germ-free larvae with *Enterobacter*-GFP at either 5 or 6 dpf. Three hours later, we fed the larvae with UV-treated rotifers (Materials and Methods). The following day (6 or 7 dpf), we imaged the larval gut with LSFM, using the green channel to visualize bacteria and the red channel to capture autofluorescence. As controls, we inoculated and imaged unfed larvae. In contrast to the large aggregates seen in unfed larvae (Fig. 2A), *Enterobacter* in fed larvae appeared planktonic and motile (Fig. 2B, Supplemental Videos 5 and 6), a behavior we had never before observed for this bacterial species. To quantify the aggregation state, we segmented objects in images of unfed and fed larvae and compared their volumes (Fig. 2C Materials and Methods). On average, the segmented objects in unfed larvae measured were well over an order of magnitude larger than in the fed larvae, with volumes 2300 ± 656 *μ*m^3^ (mean ± standard deviation) and 39 ± 5 *μ*m^3^, respectively, consistent with visual assessment of large aggregates in unfed fish and diffraction- and motion-blurred individual cells in fed fish. To quantify bacterial motility in fed larvae, we recorded videos in single optical planes at 50 frames per second and tracked *Enterobacter* cells (Fig. 2G) (Materials and Methods), finding a mean speed of 19 ± 10 *μ*m/s (Fig. 2H), consistent with the run speed observed in other *Enterobacter* strains [20].

**Figure 2.**
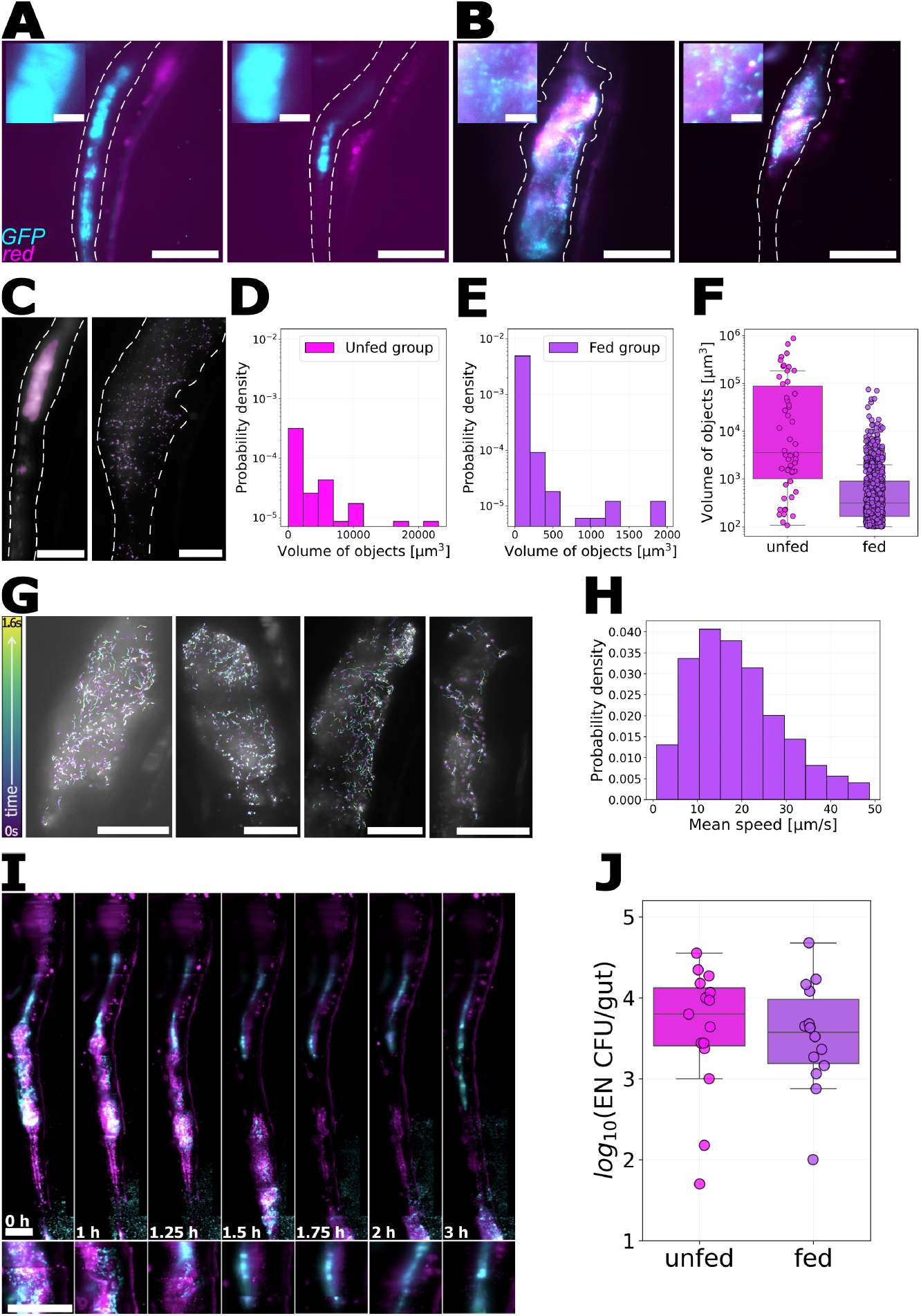
**A-B** Composite images of zebrafish intestines showing GFP-labeled *Enterobacter* (cyan) and red autofluorescence from the gut and rotifers (magenta). Dashed lines indicate approximate intestinal boundaries. Scale bar: 100 *μ*m. **A** *Enterobacter* aggregates in the midgut of an unfed larva. **B** Planktonic *Enterobacter* and rotifers in the gut of a fed larvae. Insets to (A) and (B) show zoomed-in views of the midgut; individual bacterial cells are evident in (B); scale bars: 10 *μ*m. **C** Overlay of *Enterobacter* images and their corresponding segmentation (purple) in the gut of an unfed larva (left) and a fed larva (right). Scale bar: 50 *μ*m. **D-F** Histogram of object volumes from images of **D** unfed larvae and **E** fed larvae, and **F** a summary comparison (*p* < 0.001). **G** Examples of *Enterobacter* trajectories in the gut of fed larvae. Scale bar: 50 *μ*m. **H** Probability distribution of *Enterobacter* speeds in guts of fed larvae. **I** Time-lapse of rotifer transit from anterior to the posterior gut of a larva inoculated with *Enterobacter*. After the rotifer reaches the posterior region, the bacteria again form aggregates. Images are composite overlays of the GFP signal (cyan) and autofluorescence in the red channel (magenta). Below each gut image is a zoomed-in view of the midgut, initially showing a rotifer and planktonic *Enterobacter*, followed by the appearance of bacterial aggregates.Time intervals are not uniform. Scale bar: 100 *μ*m. **J** *Enterobacter* abundance in dissected guts from unfed and fed larvae (*p* = 0.53).

Intriguingly, the behavioral shift to a planktonic, motile state appears reversible. We imaged an *Enterobacter*-inoculated, fed larval zebrafish every 15 minutes for 6 hours. After the rotifer completed its transit, *Enterobacter* re-formed into dense aggregates (Fig. 2I, Supplemental Video 7). It does not appear that rotifers alone are sufficient to induce a shift in *Enterobacter* aggregation state, however; we dissected the guts of unfed larvae containing *Enterobacter*-GFP aggregates and incubated them in petri dishes with rotifers at a concentration of roughly 1000 per ml for 3 hours (Materials and Methods). We found that the aggregates stayed intact (Supplemental Figure 2), providing evidence that the presence of rotifers being digested within the gut environment drives the bacterial transition, though it should be noted that the small volume of the intestinal lumen results in substantially greater local concentration of rotifers than is the case in the dish.

In addition to imaging, we dissected guts from fed and unfed larvae and plated them to quantify *Enterobacter* abundance (Materials and Methods). The bacterial load did not differ significantly between the two groups, with log_10_(CFU) = 3.6 ± 0.8 in the unfed group (mean ± standard deviation) and 3.52 ± 0.67 in the fed group (Fig. 2J).

Together, these findings show that rotifer ingestion changes *Enterobacter* behavior within the gut, shifting the bacterial character from aggregated and non-motile to planktonic and motile.

Next, we describe the impact of rotifer feeding on a bacterial species that behaves very differently in unfed animals: *Vibrio* ZWU0020, which is planktonic and motile in unfed larvae (Fig. 3A) [21]. We followed a protocol similar to the *Enterobacter* experiments: we began with germ-free larvae, inoculated them with *Vibrio*-GFP at either 5 or 6 dpf, fed them UV-treated rotifers three hours later, and imaged the following day (6 or 7 dpf). In parallel, we maintained a control group of unfed larvae. *Vibrio* abundance, assessed by dissection and plating of guts (Materials and Methods) was slightly lower in fed larvae (log_10_(CFU) 3.7 ± 0.7, (mean ± SD) than in unfed larvae (4.4 ± 0.4). LSFM imaging showed that in both groups *Vibrio* was planktonic and and motile in the gut (Fig. 3B, Supplemental Videos 8 and 9). To quantify this, we applied the same segmentation and tracking analysis used for *Enterobacter*. Segmented objects from both unfed and fed larvae had similar volumes: 16.9 ± 0.9 *μ*m^3^ for the unfed group (mean ± standard deviation) and 23 ± 1.6 *μ*m^3^ for the fed group (Fig. 3D–F). *Vibrio* cells showed similar swim speeds in fed and unfed fish, as assessed by tracking objects in 50 frames-per-second videos, yielding mean speeds of 34.3 ± 14.5 *μ*m/s (mean ± SD) in unfed larvae and 33.6 ± 14.3 *μ*m/s in fed larvae (Fig. 3G-I; Materials and Methods).

**Figure 3.**
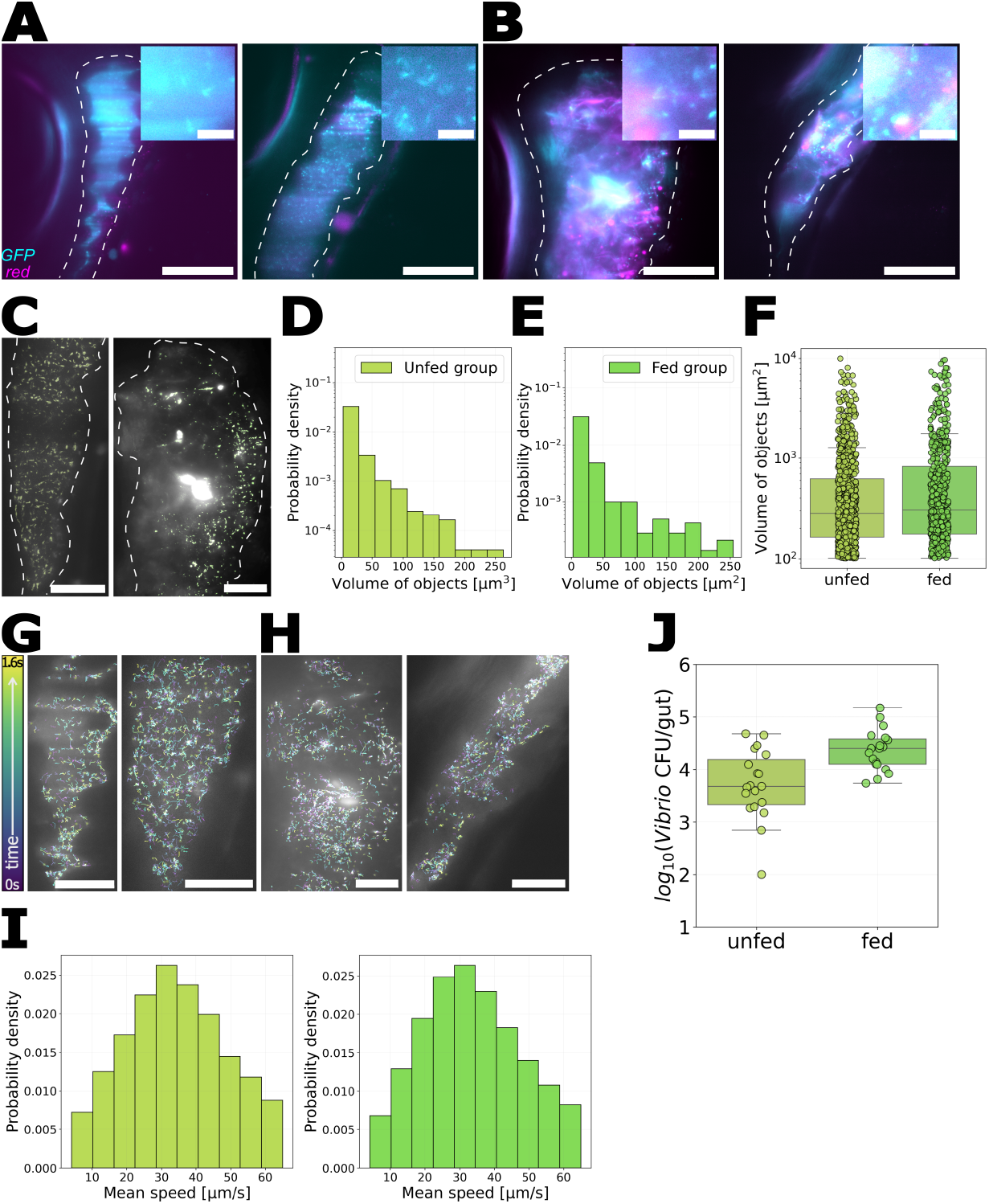
**A-B** Composite LSFM images of zebrafish intestines showing GFP-labeled *Vibrio* (cyan) and red autofluorescence from the gut and rotifers (magenta). Dashed lines indicate approximate intestinal boundaries. Scale bar: 100 *μ*m. **A** *Vibrio* in the anterior gut region of an unfed larva. **B** *Vibrio* and rotifers in the gut of a fed larva. **C** Overlay of *Vibrio* (gray) and the corresponding segmentation (green) in the guts of an unfed larva (left) and a fed larva (right). Scale bar: 50 *μ*m. **D-F** Histogram of object volumes from images of **D** unfed larvae and **E** fed larvae, and **F** a summary comparison. **G, H** Examples of tracking *Vibrio* in **G** unfed and **H** fed larvae. Trajectories are color-coded by time. Scale bar: 50 *μ*m. **I** Probability distribution of *Vibrio* speeds in guts of unfed larvae (left) and fed larvae (right). **J** *Vibrio* abundance in dissected guts from unfed and fed larvae.

As noted earlier, a key feature of the intestinal behavior of this *Vibrio* strain is the activity of its Type VI Secretion System (T6SS), a signature of which is damage to intestinal tissue, particularly resulting in widening of the gut vent (the posterior opening of the gut) [14, 15]. We confirmed prior results, finding that unfed larvae inoculated with *Vibrio* had roughly double the vent width, 103.5 ± 31.2 *μ*m (mean ± standard deviation), as germ-free unfed larvae, 51.1 ± 12.9 *μ*m, or fed larvae that were not inoculated with *Vibrio*, 50.0 ± 17.0 *μ*m. Strikingly, vent width was low in rotifer-fed larvae inoculated with *Vibrio*, with values 63.3 ± 21.1 *μ*m, closer to those of of germ-free unfed or non-inoculated fed fish than to *Vibrio*-inoculated unfed fish (Fig.4A-B). This observation led us to ask whether rotifer ingestion alters *Vibrio* behavior not by changing its organizational state or motility, but by influencing T6SS activity.

**Figure 4.**
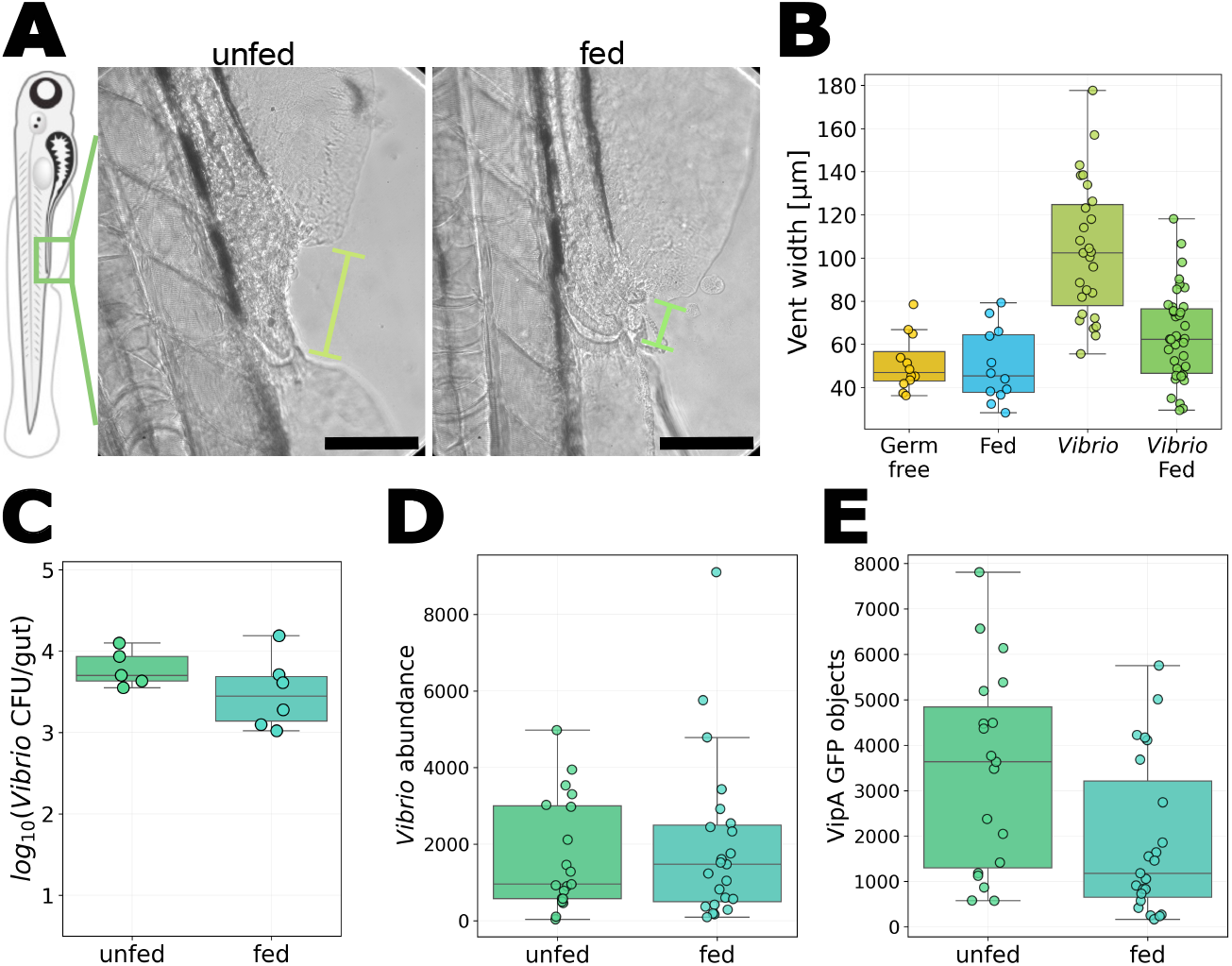
**A** Bright field images of the vent region in fed and unfed larvae mono-associated with *Vibrio*. Green lines indicate vent width. Scale bar: 100 *μ*m. **B** Vent width of larvae under various treatments: germ-free unfed, fed and not inoculated with *Vibrio*, unfed and mono-associated with *Vibrio*, and fed and mono-associated with *Vibrio*. Vent width in fed fish inoculated with *Vibrio* is roughly 60% the value in unfed *Vibrio*-inoculated fish (*p* < 0.001). **C** *Vibrio* abundance in unfed and fed larvae (*p* = 0.96). **D** Total fluorescence intensity of constitutively expressed mScarlet, proportional to overall bacterial abundance, in the vicinity of the posterior vent in unfed and fed *Vibrio*-inoculated fish. **E** The number of VipA-GFP puncta in the same fish and imaging regions as (D) (*p* = 0.015).

We repeated the *Vibrio* mono-association experiment, but this time with bacteria engineered with a reporter of T6SS activity. This *Vibrio-mScarlet;VipA-GFP* strain constitutively expresses the fluorescent protein mScarlet and encodes a fusion protein of GFP with VipA, part of the T6SS sheath assembly [22] (Materials and Methods). If imaged with high resolution [23, 24], VipA-GFP would appear as fluorescent puncta when the T6SS sheath is assembled and as diffuse fluorescence throughout the cell when VipA is expressed but the macromolecular T6SS complex is not active. Our imaging in live larvae with LSFM cannot reliably resolve subcellular puncta due to tissue-derived aberration and bacterial motion, and so VipA-GFP-positive objects may be either whole cells or T6SS assemblies. Nonetheless, the count of these objects derived using the same image analysis parameters provides a readout of relative differences in T6SS expression between experimental conditions. As in the prior experiment with *Vibrio*, assessment of bacterial abundance from plating dissected guts showed little difference between fed larvae (log_10_(CFU) = 3.5 ± 0.4, mean ± SD) and unfed larvae (3.8 ± 0.2; *p* = 0.33; Fig.4C). To determine whether feeding simply removes *Vibrio* from the vicinity of the vent, where intestinal damage is concentrated, we quantified the number of mScarlet-positive objects in the posterior quarter of the gut and found very similar numbers in unfed larvae (1708 ± 1470, mean ± standard deviation) and fed larvae (1978 ± 2150; *p* = 0.98) (Fig. 4D). To assess T6SS activity, we segmented and counted GFP-positive objects in the posterior quarter of the gut and found considerably fewer in fed larvae (1897 ± 1722, mean ± SD) than in unfed larvae (3444 ± 2196, *p* = 0.014) (Fig. 4E). Examples of images from fish with a range of VipA-GFP object counts are shown in Supplemental Figure S3. Together, these findings suggest that rotifer ingestion does not alter bacterial load or distribution, but rather alters *Vibrio* behavior in the larval gut, lowering its T6SS activity and thereby reducing damage to host tissue.

## Discussion

In this study, we used UV-treated rotifers to investigate the impact of feeding on symbiotic bacteria in the zebrafish gut. Though there is clearly a vast range of possible foods, bacterial strains, and host species, the combination of UV-treated rotifers, bacteria mono-associated with their host, and larval zebrafish provide a powerful starting point for discovery. Ultraviolet irradiation of rotifers supplies a living, nutrient-rich food source that has almost no associated microbes that would complicate analysis. The bacterial species we investigated have robust, well-defined biophysical characteristics on their own. Larval zebrafish allow control of gnotobiotic state and, crucially, direct visualization of dynamics within the intestine.

Considering a zebrafish-native *Enterobacter* strain that, in the absence of food, forms dense aggregates, we found a rotifer-induced shift to motile, planktonic cells. After the food passes, the bacteria re-aggregate. In contrast, a zebrafish native pathobiont *Vibrio* strain that is motile in the absence of food remains motile with rotifers present. However, *Vibrio* showed lower activity of its Type VI Secretion System in the intestines of fed fish than in unfed fish, leading to far less vent-associated tissue damage. The T6SS is well known as a cellular weapon, capable of delivering toxins to both bacterial and eukaryotic cells [23–25]. Intriguingly, recent work has shown that bacteria can use the T6SS to forage from their environment, lysing adjacent bacteria when starved [26]. Our observations in the zebrafish intestine suggest that a similar dynamic may be at work, with the nutrients provided by rotifer digestion satiating *Vibrio* and reducing its motivation to lyse host cells. Given the strong genetic similarity between the zebrafish-isolate *Vibrio* ZWU0020 and the human pathogen *Vibrio cholerae*, the causative agent of cholera, which also possesses a Type VI Secretion System whose activity causes strong gut contractions [15], our findings suggest a possible connection between diet and the etiology of cholera: beyond systemic host health aided by proper nutrition, food may specifically diminish *Vibrio cholerae’s* toxin deployment.

A key limitation of our work is that we have not uncovered the molecular mechanisms behind either *Enterobacter* disaggregation or *Vibrio* T6SS down-regulation. This would be worth investigating in the future through assays of gene expression and regulation, though these are more challenging to execute in a host-associated milleau than in vitro. Alternatively, specific reporters of candidate genes could be used to visualize expression within the gut environment. These approaches would help connect the behavioral changes we observe under the microscope to the underlying molecular phenomena driving them.

In this study, we focused for simplicity on mono-associations, isolating the effects of rotifer ingestion on single bacterial species. The normal state of animal gastrointestinal tracts is to house communities comprising many microbial species and examining bacterial behaviors, including specifically *Enterobacter’s* aggregation state and *Vibrio’s* T6SS activity, in a multi-species context would likely reveal a still greater richness of behavioral modulations. Similarly, expanding this work to other types of food would be interesting, especially given connections known to exist in zebrafish between food type and microbe-mediated signaling [27].

Typically, studies of the relationship between diet and the host-associated microbiome focus on changes in species abundance or community composition. Our observations here reveal that even if the number of bacteria or their persistence in the gut is unchanged, particular bacterial species can dramatically alter their behavior upon host feeding. In the case of a gut-isolate *Vibrio*, this altered behavior has direct consequences for host health. We suspect that the food-induced changes in bacterial behavior uncovered here are simply two among many that await discovery.

## Materials and Methods

### Experimental Procedures

#### Animal care

All experiments with zebrafish followed standard procedures [28] and were done in accordance with protocols approved by the University of Oregon Institutional Animal Care and Use Committee, protocol number AUP-22-02.

#### Zebrafish gnotobiology

Wild-type ABC X TU zebrafish (*Danio rerio*) were derived germfree as described previously [19] with slight modifications. In brief, fertilized eggs were collected and placed in sterile antibiotic embryo medium (EM) containing 100 *μ*g/ml ampicillin, 250 ng/ml amphotericin B, 15 *μ*g/ml gentamicin, 1 *μ*g/ml tetracycline, and 1 *μ*g/ml chloramphenicol for approximately 5 hours. The eggs were then washed in sterile EM containing 0.003% sodium hypochlorite and then in sterile EM containing 0.1% polyvinylpyrrolidone-iodine. Washed embryos were distributed into tissue culture flasks containing 15 ml of sterile embryo medium at a density of one embryo per mL. Flasks were inspected for sterility before being used in experiments.

#### Bacterial strains and inoculation of larval zebrafish

*Vibrio* ZWU0020 and *Enterobacter* ZOR0014 were previously isolated from the zebrafish intestinal tract, and fluorescently tagged strains constitutively expressing green fluorescent protein were previously generated [10]. [**ZZZ** VipA-GFP strain!] Stocks of bacteria were maintained in 25% glycerol at -80 degrees C. One day prior to fish inoculation, bacteria from frozen glycerol stocks were grown overnight in lysogeny broth (LB medium; 10 g/l NaCl, 5 g/l yeast extract, 12 g/l tryptone, and 1 g/l glucose) at 30°C with shaking. For inoculation, 1 ml of overnight culture was washed once by centrifuging for 2 minutes at 7000 × g, removing the supernatant, and adding 1 ml of fresh sterile embryo medium. Then 30 *μ*L of the bacterial suspension was added to a 15 ml tissue culture flask containing approximately 15 germ-free 5 dpf zebrafish larvae, giving a bacterial concentration of approximately 10^6^ CFU/mL.

#### Rotifers, UV irradiation, and feeding

Rotifers (*Brachionis plicatilis*) were provided by the University of Oregon Zebrafish Facility. Rotifer cultures were raised in the facility in 5 gallon containers, fed with the “Rotigrow Plus” algae mixture (Reed Mariculture), and maintained at 4 ppt salinity in “Instant Ocean” commercial sea salt mix (Instant Ocean). Rotifers at a density of roughly 2000 per liter were obtained as needed from the facility and used the same day. For ultraviolet irradiation, We follow the protocol described in [18]. In brief, we diluted rotifers 1:4 (volume/volume) with 4 ppt NaCl, and placed 15 ml of this suspension in a three-neck round bottom flask, with a stir bar, for continuous stirring. The center cap has LED which provides UV light and the remaining necks of the flask were kept loosely capped to allow oxygenation. The rotifers are exposed to 4 cycles of UV light consisting of alternating 30-minute exposure and rest periods. To feed rotifers to zebrafish larvae, we added 1 ml of the UV-treated rotifer suspension to each 15 ml flask, corresponding to roughly 400 rotifers or 30 per fish. Fish were allowed to feed ad libitum.

#### Light sheet fluorescence microscopy

Imaging was performed using a home-built light sheet fluorescence microscope based on the design of Keller et al. [29]. The protocols to mount fish and the details of the microscope can be found in references [11, 15, 21, 30]. In brief, we anesthetize larvae with MS-222 (tricaine methanesulfonate, 120 *μ*L per ml) and mount the fish in glass capillaries containing 0.7% agarose gel suspended vertically, head up, in a custom imaging chamber containing EM and MS-222. The agar-embedded larvae were partially extruded from the capillary to ensure that the optical paths did not cross glass interfaces. Lasers with excitation wavelengths of 488 and 568 nm were used to excite green fluorescent protein and mScarlet labeled bacteria, respectively. To image the entire extent of the intestine covering the gut (approximately 1200 × 300 × 150*μ*m^3^, we sequentially image four or five subregions, computationally registering the images after acquisition. Registration is performed using custom software, publicly available in a GitHub repository (https://github.com/pamitabh/batchprocessing/tree/main/use_friendly_downsampling_mip_stitch_batchprocess_code;function run2), which uses stage positions saved during image acquisition to align and stitch image stacks with the correct offsets.

#### Plating and bacterial abundance measurements

To determine the intestinal abundance of bacterial species, dissections of larval zebrafish were performed at 6 dpf or 7 dpf. Zebrafish were euthanized by hypothermal shock. Intestines were removed by dissection, placed in 1 ml sterile embryo medium, and homogenized with zirconium oxide beads using a bullet blender. The homogenized gut solution was diluted by factors of 10^−1^ and 10^−2^ and 100 *μ*l volumes of these dilutions were spread onto LB-agar plates.

#### Design and Construction of fluorescent T6SS activity reporter in Vibrio ZWU0020

The construction of a T6SS activity reporter in *Vibrio* ZWU0020 was based on a previous approach used in human-derived pathogenic *Vibrio cholerae* [24], engineering a translational fusion between the T6SS sheath protein VipA (IMG ID: ZWU0020_02771) and superfolder green fluorescent protein (sfGFP), inserted at the native vipA locus. This was done by first cloning an open reading frame encoding sfGFP, which also contained a linker peptide (3xAla 3xGly) at the N-terminus, in-frame and immediately before the stop codon of the vipA gene through a 3-piece splice by overlap extension (SOE) approach using primer pairs: WP245/WP246, WP247/WP248, and WP249/WP250. The resulting DNA fragment contained a total of 813bp and 805bp of flanking homologous DNA sequence up- and downstream of the vipA locus and was then inserted into the allelic exchange vector pAX2, with its sequenced confirmed by Sanger sequencing using the primers WP17, WP24, WP126, WP140, WP251, and WP252. The resulting vector (pTW469), was delivered by conjugation into Vibrio ZWU0020. Recombinant clones were selected and screened for as previously described [10]. Successful insertion of the vipA-sfGFP fusion gene was confirmed with PCR using primers WP251 and WP252. To fluorescently label Vibrio cells, a constitutively expressed gene encoding mScarlet was inserted into the Vibrio chromosome at the attTn7 insertion site as previously described [10]. The mScarlet construct was made by first cloning an open reading frame encoding mScarlet into the expression scaffold of pXS-sfGFP, creating pXS-mScarlet. A DNA fragment containing the constitutive promoter Ptac, mScarlet open reading frame, and trpL transcriptional terminator, was then subcloned into the Tn7 tagging vector pTn7xTS, creating pTn7xTS-mScarlet. pTn7xTS-mScarlet and the pTNS2 helper plasmid were then conjugated into Vibrio. Tagged clones were screened and selected for as previously described [10]. Tn7 insertion was confirmed by PCR using primers WP11 and WP12. Tables of strains, plasmids, and primers used for fluorescent T6SS activity reporter construction are provided as Supplemental Material.

### Image Analysis

All image processing and data analysis were performed using code written in Python using the NumPy, SciPy, scikit-image, and pandas libraries, or using Ilastik v1.4.1, as described below.

#### Autofluorescence intensities in unfed and rotifer-fed larvae without bacteria

We consider the anterior half of the gut, noting that rotifers remain longer in the anterior bulb than in the posterior gut. We use the maximum intensity projection of red channel fluorescence images to create a coarse mask that allows us to exclude the highly autofluorescent yolk from the analysis. We compute the maximum fluorescence intensity in the green channel as the maximum pixel value and the total intensity in the red channel as the sum of pixel intensities, each within the mask limits.

#### Segmentation of bacteria and bacterial aggregates

The overall gut boundary is determined by manually drawing a mask on maximum intensity projection images. Segmentation is then performed by background subtraction followed by high-pass filtering to remove low-spatial-frequency background, mainly intestinal mucus autofluorescence. The images are then thresholded and objects and their properties are found using the skimage functions ‘label’ and ‘regionprops’ to characterize connected groups of binarized voxels.

#### Assessing bacterial motility

To track bacterial motility we capture 2D videos, i.e. imaging one optical slice, at a frame rate of 50 frames per second. We subtract from each frame the median across all frames to discard objects that are not moving, which are usually parts of rotifers. We high-pass filter and mask the overall extent of the gut as above. A global threshold value across all frames is found by averaging the threshold values determined from Otsu’s method applied to each frame. As above, the ‘label’ and ‘regionprops’ functions from the skimage Python package are used to identify objects in each frame, giving a list of positions, and the ‘link_df’ function from the Trackpy package is used to link positions into trajectories. We consider 80-frame durations and remove trajectories whose maximum displacement is less than 6 pixels (1.0 *μ*m) and which last less than 5 frames for *Enterobacter*, and less than 5 pixels and 4 frames for *Vibrio*, to discard noise or non-bacterial objects, determining these parameters by direct observation. The mean distance per frame gives a mean speed value for each bacterial track. The data shown for *Enterobacter* are pooled from 8 fish; for *Vibrio* 8 fed and 8 unfed fish.

#### Segmentation of VipA-GFP objects

VipA-GFP objects were segmented using a pixel classification workflow in GPU-accelerated Ilastik v1.4.1, which employs a Random Forest classifier trained on user-annotated foreground and background labels [31]. For each imaging session, we trained separate classifiers for the bacterial constitutive reporter (mScarlet) channel and the vipA-GFP channel, retraining for data from different imaging days to account for minor variations in optical alignment. Within each channel, the same trained classifier was applied to both control and treatment groups to ensure consistent segmentation criteria. Training was performed interactively by labeling foreground (bacterial puncta or VipA objects) and background pixels across multiple z-stack slices. Default pixel features (smoothed intensity, edge filters, and texture descriptors) were computed using multi-scale Gaussian filters with sigma values ranging from σ_0_ = 0.3 to σ_6_ = 10 pixels. Prediction was monitored during training to ensure accurate segmentation, with additional labels added iteratively until classification performance was satisfactory across representative images. After training, classifiers were applied in batch mode to all images from the respective imaging session and channel. Segmentation masks were exported as 8-bit TIFF files and used for downstream quantification.

## Supporting information

Supplemental Figures, Video Captions, and Dataset Descriptions

CSV files of plotted data

## Acknowledgments

We thank Rose Sockol, Tim Mason, and the University of Oregon Zebrafish Facility staff for fish husbandry and for providing rotifers.

This work was supported by the New Tools for Advancing Model Systems in Aquatic Symbiosis program of the Gordon and Betty Moore Foundation (doi:10.37807/GBMF9320) and the National Science Foundation under award 2310570 and a Research Traineeship (NRT) Award DGE 2022168. The funders had no role in study design, data collection and analysis, decision to publish, or preparation.

## Data Availability

We provide as Supplemental Material CSV files of all plotted data. Image data consisting of three-dimensional light sheet fluorescence microscopy image stacks are available by request; each single-color single-timepoint image is roughly 10 GB in size, with the total amount of data being over 1 TB.

## Supplemental Material

Supplemental Material are uploaded for review and are also available from the following links:

- Supplemental Document, including Supplemental Figures and Supplemental Video captions: [Link](https://www.dropbox.com/scl/fi/qvezny9m1nbzun0d4jvb0/Supplemental-Material.pdf?rlkey=yma7653mk60277424o1y2vrzc&st=k5kunf7a&dl=0)
- Supplemental Videos (Folder): [Link](https://www.dropbox.com/scl/fo/3icpgxmjs9cvkkh1skgph/ALLHak-M-lZMH-WN1BcQfm0?rlkey=idam7xj5e8mvrg02qvs97zhke&st=8nj1j1at&dl=0)
- CSV files of all plotted data: [Link](https://www.dropbox.com/scl/fi/98gvp217ykou5ri4irkcb/supplementary_data.zip?rlkey=93rcmzr7qm8lu2qfibvt2p6hp&st=bgzqcxv4&dl=0)

## Conflicts of Interest

The authors declare no conflicts of interest.

## Author Contributions

Susana Márquez Rosales: Conceptualization, Methodology, Software, Formal Analysis, Investigation, Writing – Original Draft Preparation, Visualization. Piyush Amitabh: Conceptualization, Methodology, Software, Formal Analysis, Investigation, Writing – Review & Editing, Visualization. Emily M. Olmstead: Investigation. Emily P. R. Avey: Investigation. Elena S. Wall: Resources. Lizett Ortiz de Ora: Resources. Travis J. Wiles: Resources, Review & Editing, Supervision. Raghuveer Parthasarathy: Conceptualization, Methodology, Formal Analysis, Investigation, Writing – Review & Editing, Supervision, Project Administration, Funding Acquisition.

## Notes

### Competing Interest Statement

The authors have declared no competing interest.

### Summary of Updates

Minor changes to the abstract, Introduction (paragraph 2), and the concluding discussion. No changes to data or figures.

