## Supplemental Figures, Video Captions, and Dataset Descriptions for "Imaging the impact of rotifer consumption on bacterial behaviors in the zebrafish gut"

#### Contents

- Description of Supplemental Data
- Supplemental Movie Captions
- Supplemental Figures and Captions
- Tables of strains, plasmids, and primers used for fluorescent T6SS activity reporter construction

#### Description of Supplemental Data

The following are contained in a compressed (ZIP) file, supplementary_data.zip:

**Plotted Datapoints.** CSV files of all plotted datapoints, organized by figure.

### Supplemental Movie Captions

**Supplemental Movie 1.** Animated z-stack of light sheet fluorescence microscopy images of unfed larval anterior gut and surrounding regions, shown in Figure 1B. Cyan: green autofluorescence, magenta: red autofluorescence as explained in the main text.

**Supplemental Movie 2.** Animated z-stack of light sheet fluorescence microscopy images of fed larval anterior gut and rotifers, shown in Figure 1C. Cyan: green autofluorescence, magenta: red autofluorescence as explained in the main text.

**Supplemental Movie 3.** A series of maximum intensity projections of 3D images taken from time lapse, every 15 minutes, imaging rotifers transit within a fed larval zebrafish gut (Figure 1E). Composite overlays: cyan shows green autofluorescence and magenta shows red autofluorescence, shown in Figure 1E.

**Supplemental Movie 4.** Animated z-stack of light sheet fluorescence microscopy images of unfed zebrafish intestines (6 dpf) showing GFP-labeled Enterobacter (EN) (cyan) and red autofluorescence from the gut (magenta), shown in Figure 2A.

**Supplemental Movie 5.** Animated z-stack of light sheet fluorescence microscopy images of fed zebrafish intestines (6 dpf) showing GFP-labeled *Enterobacter* (cyan) and red autofluorescence from the rotifers (magenta), shown in Figure 2B.

**Supplemental Movie 6.** Light sheet fluorescence microscopy movie of motile *Enterobacter* within a fed larva. Each frame is from the same optical plane, which spans the midgut region (Fig. 2B).

**Supplemental Movie 7.** A series of maximum intensity projections of 3D images taken from time lapse, every 15 minutes, imaging GFP-labeled *Enterobacter* (cyan) and rotifers transit (autofluorescence in magenta) within a fed larval zebrafish gut (Figure 2I). After rotifers reach the vent, *Enterobacter* forms aggregates again.

**Supplemental Movie 8.** Animated z-stack of light sheet fluorescence microscopy images of unfed zebrafish intestines (6 dpf) showing GFP-labeled *Vibrio* (cyan) and red autofluorescence from the gut (magenta), shown in Figure 3A.

**Supplemental Movie 9.** Animated z-stack of light sheet fluorescence microscopy images of fed zebrafish intestines (6 dpf) showing GFP-labeled *Vibrio* (cyan) and red autofluorescence from the rotifers (magenta), shown in Figure 3B.

**Supplemental Movie 10.** Light sheet fluorescence microscopy movie of motile *Vibrio* in an unfed larva. Each frame is from the same optical plane, which spans the midgut region (Fig. 3A).

**Supplemental Movie 11.** Light sheet fluorescence microscopy movie of motile *Vibrio* within a fed larva. Each frame is from the same optical plane, which spans the midgut region (Fig. 3B). The most bright non-motile regions correspond to ingested rotifers.

### Supplemental Figures

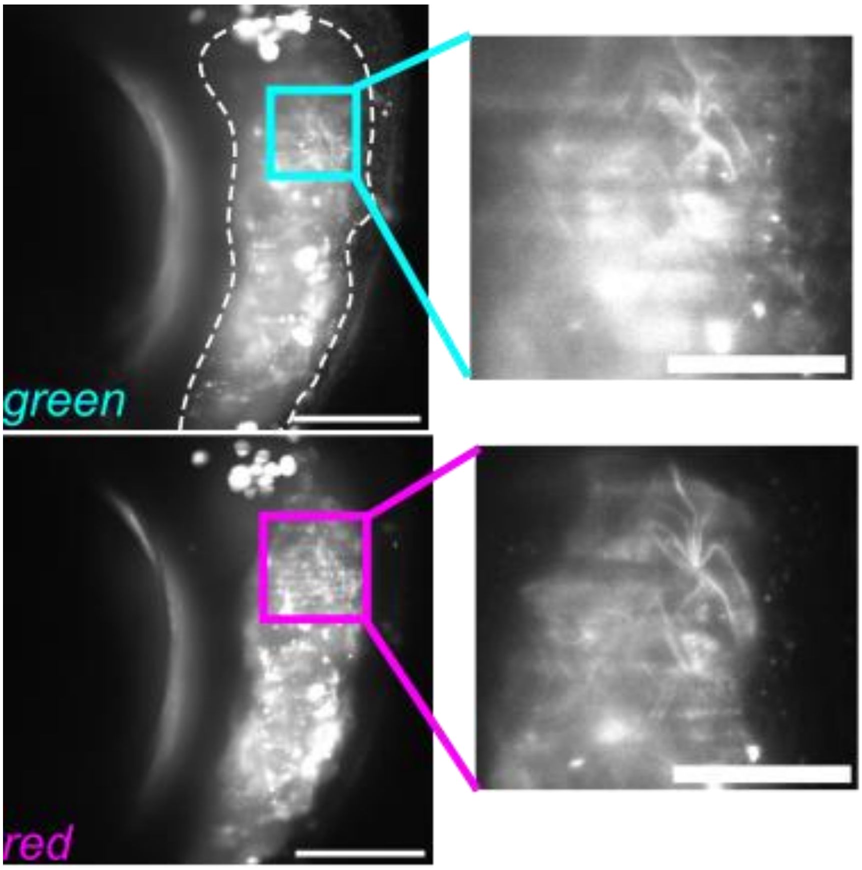


**Supplemental Figure S1.** Anterior gut of fed zebrafish autofluorescence imaged with light-sheet fluorescence microscopy (LSFM) using blue/green excitation/emission wavelengths (excitation 488 nm, emission 510-550 nm, denoted 'green') and autofluorescence using green/red excitation/emission wavelengths (excitation 561 nm, emission 590-650 nm, denoted 'red').

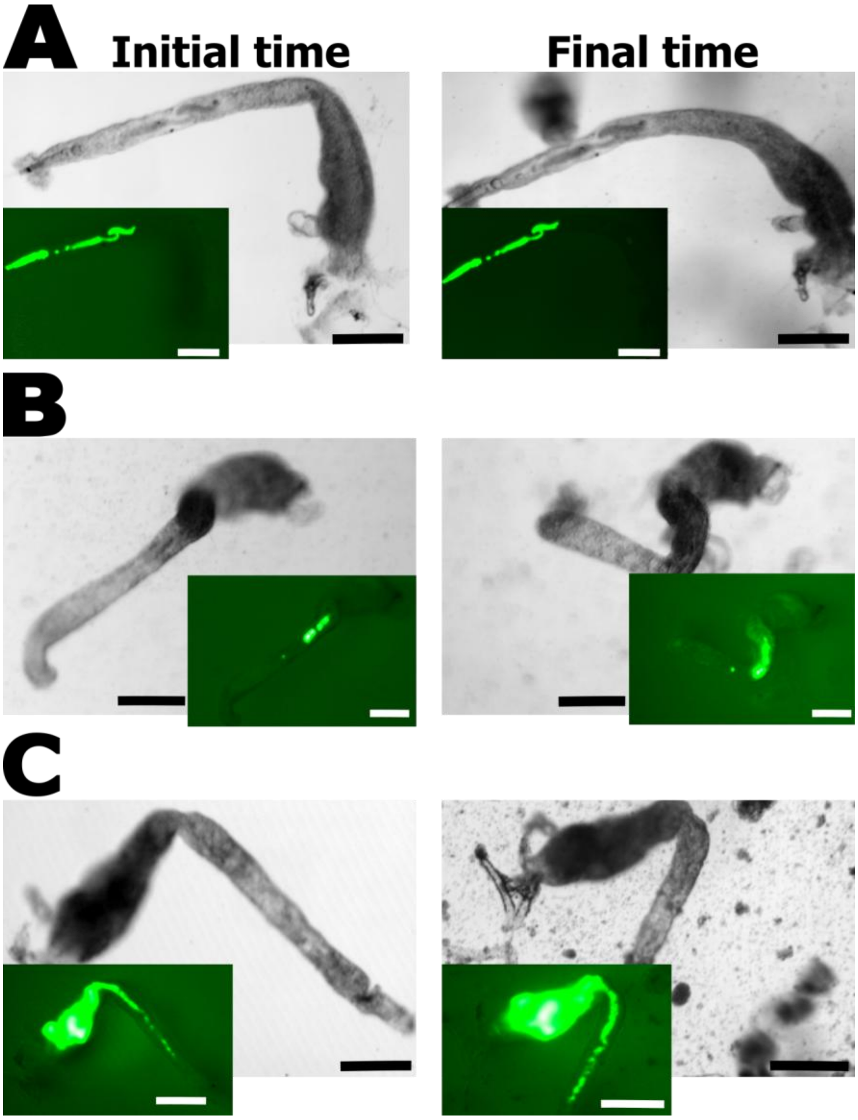


**Supplemental Figure S2.** Dissected guts of zebrafish larvae containing fluorescent EN aggregates. Panels A–C correspond to different larvae. Left: Initial time, right: final time. The gray picture shows the bright field image and the green images show the *Enterobacter* GFP. Scale bar: 200 μm.

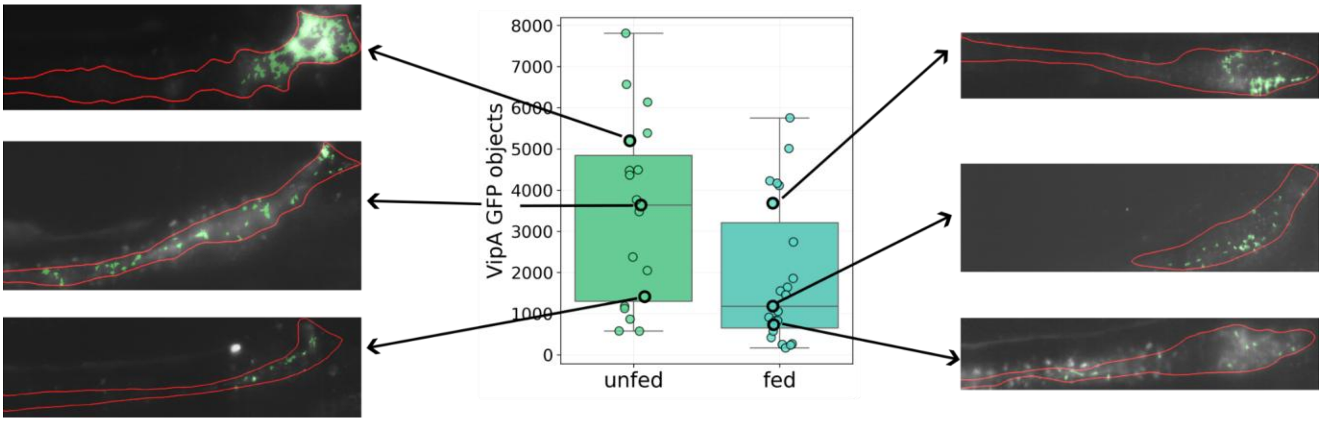


**Supplemental Figure S3** Examples of segmented VipA-GFP objects in unfed and fed larvae (see Figure 4E). The red line outlines the gut mask; green regions are segmented objects.

### Tables of strains, plasmids, and primers used for fluorescent T6SS activity reporter construction

##### Strains

| Strain | Description/ Relevant Details | Reference/Source |
|---|---|---|
| DH5α | general cloning strain | NEB |
| SM10(λpir) | pir⁺ mating strain; $Kan^R$ | REF |
| *Vibrio* ZWU0020 | *Vibrio cholerae* strain isolated healthy zebrafish gut | |

##### Plasmids.

| Plasmid | Description/ Relevant Details | Reference/Source |
|---|---|---|
| pAX2 | allelic exchange vector with *GFP*$^{mut3.1}$ merodiploid tracker, temperature-sensitive replicon *ori*$_{101}$/*repA101*$^{ts}$, and TetR-controlled kill switch; $Amp^R$, $Gent^R$, $Clm^R$, 30°C | Wiles et al, 2018 |
| pTW469 | pAX2 with an allelic exchange cassette containing the *vipA-sfGFP* fusion; $Amp^R$, $Gent^R$, $Clm^R$, 30°C (Strain # TW469) | This study (or Cathy's?) |
| pmScarlet_C1 | pC1 vector with monomeric (m)Scarlet template; obtained as a gift from Dorus Gadella (Addgene plasmid # 85042; http://n2t.net/addgene:85042; RRID:Addgene_85042) | REF |
| pXS-sfGFP | Vector with a modular sfGFP expression scaffold; $Amp^R$ | Wiles et al, 2018 |
| pXS-mScarlet | pXS vector with constitutive mScarlet gene (Strain # pTTW3) | |
| pTn7xTS | Tn*7* tagging vector with temperature-sensitive replicon *ori*$_{101}$/*repA101*$^{ts}$; $Amp^R$, $Gent^R$, 30°C | Wiles et al, 2018 |
| pTn7xTS-mScarlet | Tn*7* tagging vector pTn7xTS temperature-sensitive replicon *ori*$_{101}$/*repA101*$^{ts}$; $Amp^R$, $Gent^R$, 30°C (Strain # TTW8) | This study |
| pTNS2 | pTNS2; Tn*7* helper plasmid carrying Tn*7* transposase genes; $Amp^R$ | REF |

##### Primer sequences.

| Name | Sequence (5'–3') |
|---|---|

| | |
|---|---|
| seq.pUC18R6KT-Tn7T-T1-REV (WP17) | cttaaacgcctggggtaatg |
| hokB.Ptac_SOE.FOR (WP24) | tgagcggataacaatttcacacaggagaaaggctatgaagcac |
| SFGFP.REV (WP126) | tcacttgtacagctcgtccatg |
| seq.sfGFP.5p.REV (WP140) | ctgaacttgtggccgtttac |
| 5p.HR.SmaI.vipA.FOR (WP245) | cccgggtattagatcatcgcaaaca |
| 5p.HR.vipA.3A3G.sfGFP.SOE.REV (WP246) | agctcctcgcccttgctcataccaccgcccgcagctgccgcttgtggctct tcttgac |
| 3A3G.sfGFP.SOE.vipA.FOR (WP247) | gtcaagaagagccacaagcggcagctgcgggcggtggtatgagcaagggcg aggagct |
| sfGFP.SOE.vipA.REV (WP248) | gtgattaattgattaataacgtttgctcacttgtacagctcgtccatg |
| 3p.HR.sfGFP.vipA.SOE.FOR (WP249) | catggacgagctgtacaagtgagcaaacgttattaatcaattaatcac |
| 3p.HR.SmaI.vipA.REV (WP250) | cccgggtggattttcgattggatcgt |
| seq.vipA.FOR (WP251) | ccgagcttcctgttgaactc |
| seq.vipA.3p.HR.REV (WP252) | tgcagcagcttcatatctgg |
| mScarlet.IVA.pXS.F | tttgtttaactttaagaaggagatatacatatggtgagcaagggcgag |
| mScarlet.IVA.pXS.R | tttgtagagctcatccatgccatgtgtgcggccgcttacttgtacagctcg tccatgcc |
| Tn7R.PCRver.FOR (WP11) | cacgcccctctttaatacga |
| Tn7.insert.ZWU0020.2.FOR (WP12) | agggtaccgatgttgaccag |
